# Reduced size of larvae and small fish linked to warming and reduced prey density

**DOI:** 10.1101/2025.06.27.661664

**Authors:** Max Lindmark, Malin Werner, Peter Thor, Federico Maioli, Eros Quesada, Valerio Bartolino, Philip Jacobson

## Abstract

Body size is a integrative trait that has declined across ecological assemblages in the last decades. Changes in body size of early life stages and small-bodied fish are driven by complex interactions between size-dependent mortality, and temperature- and food dependent growth, and can have consequences for recruitment to the adult stock and food availability to predators. While the mechanisms can be difficult to disentangle using observational data alone, simple indicators such as mean size can provide important information about ecosystem conditions as oceans rapidly change. Using 30 years of observations from the IBTS-MIK ichthyoplankton survey in Skagerrak and Kattegat in combination with sea surface temperature data, we investigated changes in the body size of early life stages and small-bodied fishes for 10 commercial and non-commercial species using geostatistical mixed models, and trends in prey density using generalized additive models. We found positive associations between chlorophyll-a concentration and length in 8 species (3 significant), and negative associations between temperature and length in 9 species (4 significant). Standardized indices of length revealed negative trends over time for all species since 2010, and all species were smaller in 2024 compared to 2000. The decline in size since 2010 varied between 30%–1%, with a mean of 14% and 9% for species predominantly found in Skagerrak and Kattegat, respectively. The trend of decreased body sizes since 2010 coincides with rapid declines in *Calanus* spp. — a key prey predicted to decline in this area at the edge of their distribution area due to climate change. In Skagerrak it also coincides with a decline in the density of large copepods (*>* 0.25 mm). The synchronous declines in larval size across taxonomically diverse species experiencing different rates of exploitation suggest a common response to changing environmental conditions, which could have cascading effects throughout marine food webs.

## Introduction

Over the past century, significant changes in the body size of organisms have been observed across a wide range of species and assemblages (Martins *et al*. 2023). These patterns tend to be stronger in marine fish, and have been attributed to size selective exploitation (Bosch *et al*. 2022) and climate change (Daufresne *et al*. 2009). Climate change can affect organism body size via multiple interlinked processes. For example, warming leads to elevated metabolic rates, and unless warming is avoided via behavioral changes or range shifts, this increase need to be met by increased feeding rates to maintain similar growth (Brett *et al*. 1969, Huey and Kingsolver 2019). On a population and community level, warming can also affect the size structure of populations and communities by increasing mortality rates (Lorenzen 2022, Lindmark *et al*. 2023b). Body size is a fundamental trait to many aspects of biology and ecology, influencing metabolism, predator-prey interactions, and ultimately affecting higher levels of biological organization (Elton 1926, Brown *et al*. 2004, Andersen 2019). Therefore it is important to quantify changes in body size across a range of species, life histories, and life stages.

Larvae, with their small body size, high metabolic rates, limited energy reserves, and low swimming capacity, are thought to have narrower ranges of tolerable environmental conditions than larger conspecifics (Irvin 1974, Brewer 1976, Rijnsdorp *et al*. 2009, Pörtner and Peck 2010). This makes them more vulnerable to changes in environmental conditions, such as ocean warming. Moreover, several studies have suggested fish larvae being food limited (Ware and Lambert 1985, Buckley and Lough 1987, Anderson 1988, Kiørboe *et al*. 1988), meaning that unless warming leads to proportional increases in food availability, growth could be expected to decline (Rogers *et al*. 2021). This could affect biomass and abundance negatively through lower survival rates. Yet, despite the higher sensitivity to climate change and changes in feeding conditions, our understanding of these effects on early life stages remains limited. This is especially true in natural environments where data are often sparse or difficult to collect compared to commercial fish of landing size (Leggett and Deblois 1994).

In fish, recruitment to the adult life stage of the population is largely determined by processes occurring during these early stages (Hjort 1914, 1926, Ricker 1954, Myers 1995). Hence, understanding the impacts of climate change on the size of fish larvae is important for predicting the impacts on population dynamics (Rijnsdorp *et al*. 2009). There are numerous hypotheses about the processes that affect larval survival and thereby recruitment, and many of them are related to growth and body size. For example, the “match-mismatch hypothesis” (Cushing 1973, 1990), states that overlap between larvae and plankton production shape the survival of early life stages (by avoiding starving and increasing growth) and ultimately determines the recruitment strength of temperate fishes. Faster growth allows larvae to spend less time in the high-mortality pelagic stage, and thereby experience a lower cumulative mortality risk, since size rather than age determines metamorphosis (Chambers *et al*. 1988). This mechanism is also central in the “stage-duration hypothesis” (Leggett and Deblois 1994), which can be viewed within a broader “growth-mortality hypothesis” (Werner and Gilliam 1984, Anderson 1988). Research accumulated over the last century has shown that pre-recruit survival is not determined solely by any of these hypotheses. Partly this may be due to a combination of these difficult-to-test hypotheses and a lack of adequate data. However, evidence suggests that larvae survival and thereby recruitment is influenced by the interrelated processes of feeding, body growth and predation — all of which are likely to be affected by warming oceans (Leggett and Deblois 1994).

Given the sensitivity of early life stages to environmental fluctuations, and the role of growth and size in affecting survival of fish, studying changes in body size over time can serve as a valuable indicator of how environmental conditions for early life stages are shifting. Research on larval fish responses to climate change has historically focused on commercially important species. However, non-commercial species are as important for ecosystem functioning, and there is a risk of confounding climate impacts with those of commercial exploitation if only focusing on commercial species. This because of the effects of fishing on the demography of stocks (Barnett *et al*. 2017, Griffiths *et al*. 2024), which can have effects on spawning phenology and thereby larvae size at date (Rogers and Dougherty 2019), or on the size and quality of eggs (Hixon *et al*. 2014). Fishing may also have synergistic effects with climate on body growth and developmental rates (Audzijonyte *et al*. 2016). This highlights the importance of evaluating climate impacts on species from diverse taxonomies, body size range, and different exploitation histories. Lastly, most studies of body size changes do not account for spatial dynamics (but see Indivero *et al*. 2023), including range shifts and interannual variation in spatial distribution, which can influence observed body size trends.

In this study, we apply geostatistical models to data on body size of 10 larvae or small bodied species, commercial and non-commercial (e.g., gobies, pipefishes, pricklebacks, gunnels, and sculpins) from three decades of ichthyoplankton surveys in the Skagerrak and Kattegat. Specifically, we aim to: 1) Quantify effects of local temperature and chlorophyll-a concentrations on body sizes; 2) Estimate trends in standardized indices of length and relative biomass between 1993–2024 from spatiotemporal models; 3) Evaluate the association between average trends in body size with time series of the key copepod prey *Calanus* spp., and an index of small copepod density.

## Methods

### Fish data

Fishes were sampled in Kattegat and Skagerrak during the International Bottom Trawl Survey between 1993–2022. The survey is typically conducted over a three-week period, from late January to mid-February. Fish larvae and small bodied fish were sampled during the period between 30 minutes past sunset to 30 minutes before sunrise (ICES 2013). The main target of the survey is the larvae of autumn-spawning herring (*Clupea harengus*), but over-wintering larvae of other species and small bodies fish species are also caught (ICES 2013, Munk *et al*. 2014).

Between 2–5 locations were selected in each ICES standard rectangle (1°longitude × 0.5°latitude) (Fig. 1). At each location, hauls at a speed of 3 knots were conducted using a 2-meter diameter Midwater Ringnet (MIK), which was equipped with a 13 meter long black netting (1.6 mm mesh size) and two-legged bridles attached to both the hauling wire and depressor. A flowmeter was mounted at the nets opening to measure the volume of water entering the net. The gear was towed behind the ship going down to a minimum depth of 5 meters above the seabed or to a maximum depth of 100 meters. Depth measurements were recorded using a gear-mounted depth sounder, while the bottom depth was estimated by comparing readings from the ship-mounted echo sounder. The net was fitted with a fine-meshed hind part (0.5 mm mesh size) and a similar mesh in the cod-end bucket (ICES 2013).

**Figure 1:**
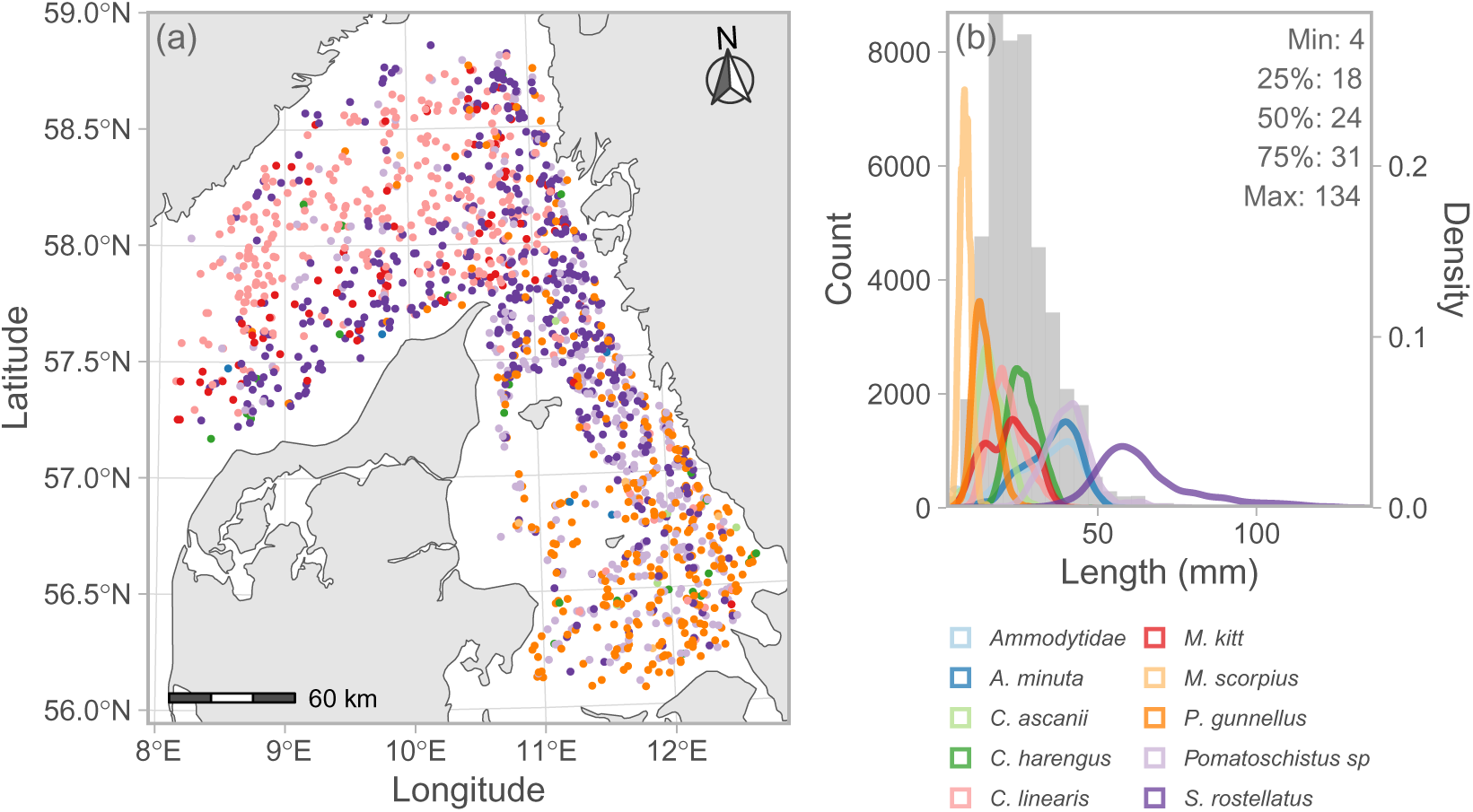
Location of hauls, where color denotes species (a), and the size-distribution of larvae and small-bodied fish used in the length models (b). ICES rectangles are depicted in panel a.

After each haul, the hind part and cod-end were flushed, and the collected plankton and fish were preserved in either 4% formaldehyde-freshwater solution or in 70% ethanol. Fish larvae and juveniles were identified to the lowest possible taxonomic level, and their standard lengths (for clupeid and gadoid species) or total lengths (for other species) were measured to the nearest millimeter.

Fish density (numbers m^-2^) were calculated as:

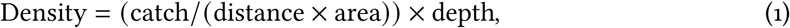

where density is numbers per m^2^, catch is number of caught, distance is the towed distance (m) as measured by the calibrated flowmeter, area is the opening of the ring (m^2^), and depth is the water depth (m). The variable density therefore represents the abundance of larvae and small fish below a certain surface area (ICES 1993, Munk *et al*. 2014). The MIK is assumed to have a catchability of 100% and is used to calculate indices of larvae abundance by expanding densities to the entire spatial domain (ICES 1993, 2013).

While all specimens are measured for length and density, we proceed with species that had at least eight individuals in a given year, and at least a time series of 18 years. The survey is an ichthyoplankton survey, but for some species also non-planktonic stages where recorded and their inclusion varied over time. For consistency and to avoid potential bias due to inclusion of sporadically caught adults, we defined a size threshold corresponding to the post-larvae size for species predominantly caught as larvae, after which metamorphosis begins. Values for these sizes were taken from literature (Rusell 1976). For the small bodied species that are routinely caught in this survey, we did not implement a size threshold and used all sizes, including adults. This resulted in a dataset of 45,613 length observations across 10 species, and 1,601 hauls with spatially referenced information on larvae density (Fig. 1).

### Spatiotemporal modelling

To quantify temporal trends in individual length and relative abundance, we fit spatial and spatiotemporal (respectively) generalized linear mixed models (GLMMs) using the The Stochastic Partial Differential Equation (SPDE) approach.

For the length model, we used a generalized gamma observation model (Dunic *et al*. 2025) with a log link because fish individual lengths are positive and tend to be right-skewed. This is a flexible model that becomes a lognormal if the “Generalized gamma Q” approaches zero and gamma if Q matches the dispersion parameter (Dunic *et al*. 2025), in the Prentice (1974) parametrization. The model included independent intercepts for each year (Thorson *et al*. 2015, Thorson 2019), and linear effects of chlorophyll-a, temperature, and day of the year to correct for variability in the sampling date. Chlorophyll-a concentrations were used as a proxy for the spatial and temporal availability of zooplankton prey. This was necessary due to the scarcity of spatial data for zooplankton in the area (but see below). Chlorophyll-a and temperature at 5 meters depth were retrieved from an ensemble of statistically downscaled CMIP6 models for the climate scenario SSP2-4.5 (Kristiansen and Butenschön 2022, Kristiansen *et al*. 2024). We used median ensemble values, all covariates where scaled by subtracting their mean and dividing by their standard deviation. We also included spatiotemporal random effects to model spatial dependency in residuals, included as Gaussian Markov random fields (GMRFs). These represent spatially structured latent processes that are independent for each year (Thorson *et al*. 2015).

Similar spatiotemporal models were fitted to larvae density to 1) quantify trends in relative abundance, and 2) to use as weights when estimating annual indices of larvae length. The latter is needed since larvae are not homogeneously distributed in space, and their distribution may vary interannually and directionally over time (Indivero *et al*. 2023, Lindmark *et al*. 2023a). We modelled larval density (numbers m^-2^) with a Tweedie distribution (Tweedie 1984, Anderson *et al*. 2024) and a log link. To avoid using the same covariates as in the length model, we included independent intercepts for year, and spatial and spatiotemporal random effects using GMRFs. The spatial random effects here represent latent spatial processes that are constant across all years, representing static oceanographic covariates.

The spatial and spatiotemporal models fitted to each species separately can be written as:

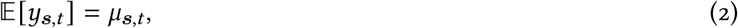

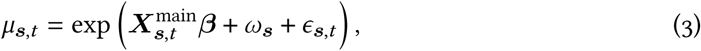

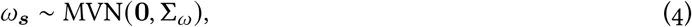

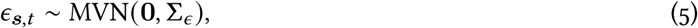

where *Y****_s_***_,*t*_ is the response variable (individual length in mm or density in numbers per m^2^) in location ***s*** (a vector of two UTM zone 32N coordinates) at time *t*, *μ* is the mean, ***X***^main^ is the design matrix for fixed effects with corresponding coefficients *β*. Spatial and spatiotemporal random effects (*ω****_s_***, *ε****_s_***_,*t*_), are assumed drawn from GMRFs with covariance matrices (inverse precision matrix) (**Σ**_*ω*_, **Σ**_*ε*_) constrained by the Matérn covariance function (Rue *et al*. 2009).

The SPDE approach (Lindgren *et al*. 2011) involves triangulation of the domain (a mesh) for calculating the three sparse matrices needed for calculating the precision matrix. We defined this mesh using triangles with a minimum distance between vertices (cutoff) of 8 km and 4 km for the length and the density models, respectively. We ensured that the estimated range, i.e., the distance where the correlation between two points effectively disappears (Lindgren *et al*., 2011), was larger than 2 times the cutoff (mean range was 38 km for the length model and 62 km for the density model). The mesh was constructed using the R-function `fm_rcdt_2d_inla()` from the package `fmesher` (Lindgren *et al*., 2023) at their defaults (see Supporting Information Fig. S1 for an example).

To evaluate trends in over time for each species, we predicted our length and density models on a 3×3 km UTM grid covering the survey domain for all years when the species was observed. From these predictions, bias-corrected, annual indices were acquired using the function `get_index()` in the R-package `sdmTMB` (Thorson and Kristensen 2016, Anderson *et al*. 2024). In the length model, we used predicted larvae density as a weight when calculating the average to account for spatial heterogeneity in length and range shifts. For the abundance indices, we used the area of the grid cell as weights. We also calculated center of gravity and their confidence intervals from the spatiotemporal models to assign main ocean area to fishes (Supporting Information Fig. S2).

We fit the models in R version 4.3.2 (R Core Team 2024), using the package `sdmTMB` (Anderson *et al*. 2024), version 0.6.0.9034. The R package `sdmTMB` implements maximum marginal likelihood estimation from the R package `TMB` (Kristensen *et al*. 2016), and sets up sparse matrices using the R package `fmesher` (Lindgren 2023). We evaluated model convergence by confirming that the Hessian matrix was positive definite, and the maximum absolute log-likelihood gradient with respect to fixed effects was *<* 0.001. We calculated simulation-based quantile residuals (Dunn and Smyth 1996, Gelman and Hill 2006, Hartig 2022, Anderson *et al*. 2024), with random effects taken from a single draw from their multivariate normal distribution (Waagepetersen 2006, Thorson and Kristensen 2024) to visually inspect consistency between data and the models (Supporting Information Figs. S3– S4).

### Association between fish larvae size and the copepod prey

Data on copepod abundance density (numbers m^−3^) are available at a monthly resolution, but not for the entire time period of the fish larvae data, and only from a few locations in Skagerrak and Kattegat. Therefore, we opted to compare time series of copepod prey density with the time series on larvae lengths. Copepod indices were estimated using data on copepodites and adults of *Calanus finmarchicus* and large copepods (*>*0.25 mm). Since there is no discrimination in the data set between *C. finmarchicus* and *C. helgolandicus*, a sibling species which also is abundant in these waters (Bonnet *et al*. 2005), we used the genus form *Calanus* spp. *Calanus* spp. are lipid-rich key prey that are vital to many of the planktivorous fish species and early life stages of fish, especially in winter (Last 1989) when our fish samples are taken. However it is not the only prey fish feed on, and they are on the larger side of prey for the fishes analyzed here. Typical *Calanus* size are between 2–4 mm. Assuming that larvae have an optimal predator-to-prey length ratio of 5% (as found in herring Hauss *et al*. 2023), and that our fish are between 4–134 mm (Fig. 1) with a median of 24, the optimal prey size for our fish range between 0.2–6.7 mm. To also include other prey species and a slightly smaller sizes of prey, we also used data on the density of large copepods (*>*0.25 mm).

We used data from the Å17 station in the Skagerrak (2007–2002) and the Anholt E station in the Kattegat (1996–2022) because they were nearest the center of gravity of the fishes and had the longest running data series (Fig. 2, Supporting Information Fig S2). Data on copepod density were downloaded from the Swedish Oceanographic Data Centre (Shark) in March 2024. At sampling events where no data are available for a specific species, we assumed zero density for that particular species at that particular time. Copepod biomass density were calculated from abundance desnity using weight/length regressions for each species (Mauchline 1998).

**Figure 2:**
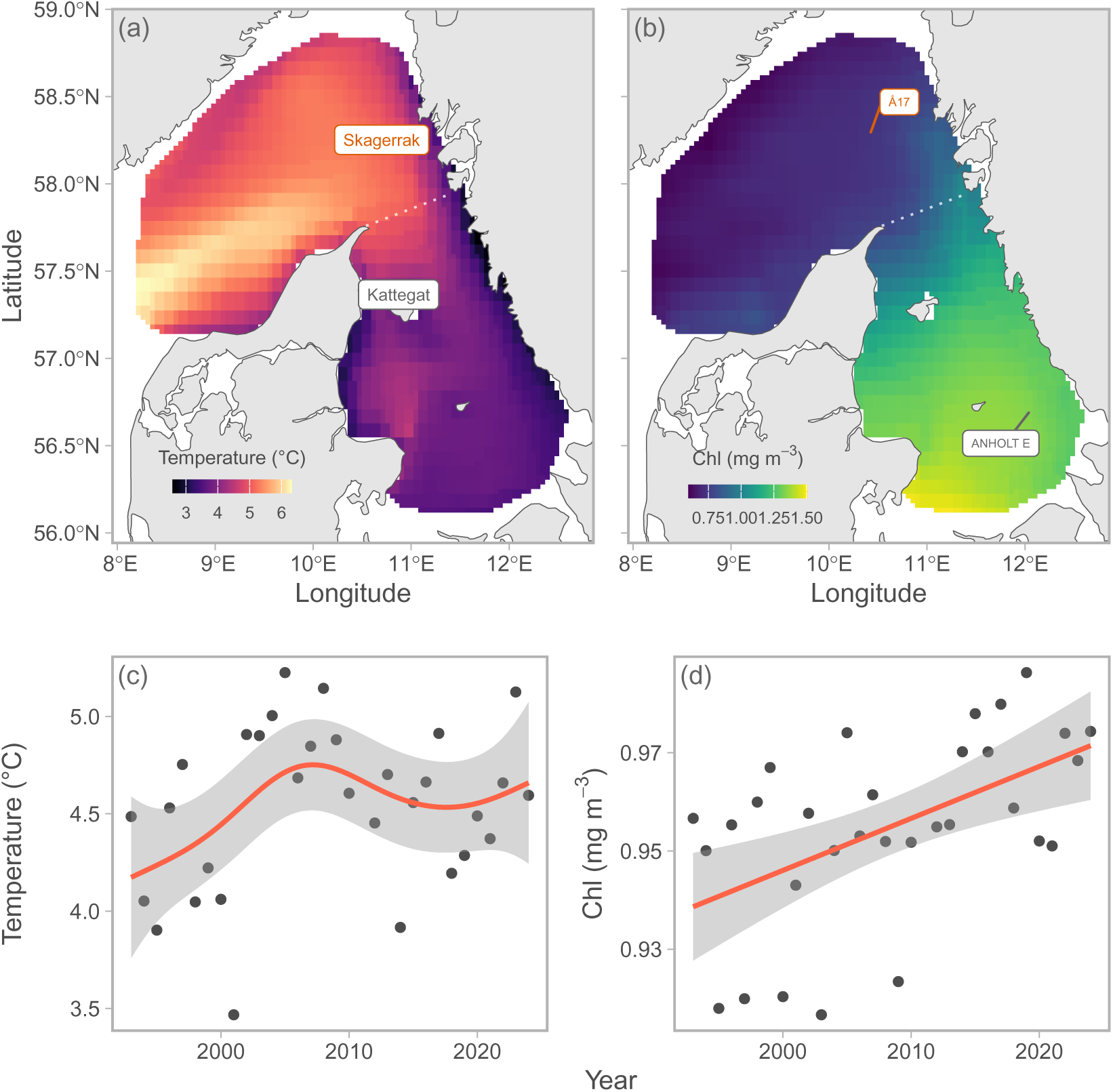
Temporally averaged sea surface temperature (a), chlorophyll-a concentration in the spatial domain (b), spatially averaged temporal trends in sea surface temperature (c), and chlorophyll-a (d) for years 1993–2024. The Skagerrak-Kattegat border is depicted in panel a. Stations for copepod data are shown in panel (b).

We standardized time series of *Calanus* spp. and large copepod density by fitting Generalized Additive Models (GAM). We modelled log densities with a Normal distribution:

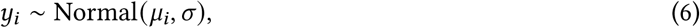

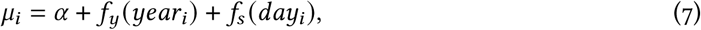

where *y*_*i*_ is log density, *μ* is the mean and *σ* is the standard deviation, *f*_*y*_ is a thin plate smoother that represents long term trends, and *f*_*s*_ represents seasonal trends. We used a cyclic cubic spline for *f*_*s*_, to avoid discontinuity at the end points of the spline (Dec 31^st^ and January 1^st^).

To acquire average trends in length across species, we fitted hierarchical GAMs (Pedersen *et al*. 2019) to *z*-scored annual length indices for all species from the spatiotemporal models, and extracted the global prediction. We modelled these annual indices separately by ocean area, with a Normal distribution:

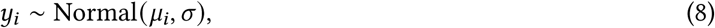

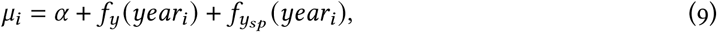

where *y*_*i*_ is length, *μ* is the mean and *σ* is the standard deviation, *f*_*y*_ is a thin plate smoother that represents long term global trend, and *f*_*ysp*_ is a factor-smoother that captures species-specific variation in trends that are shrunk towards 0. This formulation also incorporates species specific intercepts as a part of the penalty construction (Pedersen *et al*. 2019). Similarly as for the spatiotemporal models, distributional assumptions were assessed by visually inspecting simulation-based quantile residuals (Dunn and Smyth 1996, Gelman and Hill 2006, Hartig 2022) (Supporting Information Figs. S5, S6).

## Results

Over the last 30 years, the average winter temperature showed a clear pattern with the coolest temperatures in the coastal areas of Kattegat, and the warmest temperatures in the western and northwestern Skagerrak (Fig. 2a). Within the survey domain, temperatures range from 2–7 °C, and the average sea surface temperature increased from approximately 4.2 to 4.6 °C over time (Fig. 2c). Chlorophyll-a concentration showed an almost opposite pattern, with the highest concentrations in the southwest and lowest concentrations in the northwest (Fig. 2b). Over time, January chlorophyll concentration increased from approximately 0.94 to 0.97 mg m^−3^ (Fig. 2d).

The length of all species with a larvae stage showed negative associations with temperature, of which two did not have confidence intervals overlapping zero (Fig. 3a). In the small-bodied fishes, the length of three species (*Aphia minuta*, *C. linearis*, and *S. rostellatus*) was negatively associated with temperature, but the 95% confidence interval of the latter overlapped zero. *Pomatoschistus* spp. showed a weak non-significant association (Fig. 3b). The effect of chlorophyll-a concentration varied across species and life stages. The length of all larvae was positively associated with chlorophyll-a (Fig. 3c), but confidence intervals overlapped 0 for *Ammodytidae*, *C. ascanii*, and *M. scorpious*. Of the small-bodied fish, most effects were near zero, but in *S. rostellatus* a clear negative association was found (Fig. 3d). Day of the year had a positive effect on nine of ten species, but all effects were very minor compared to those of temperature and chlorophyll (Fig. 3e,f).

**Figure 3:**
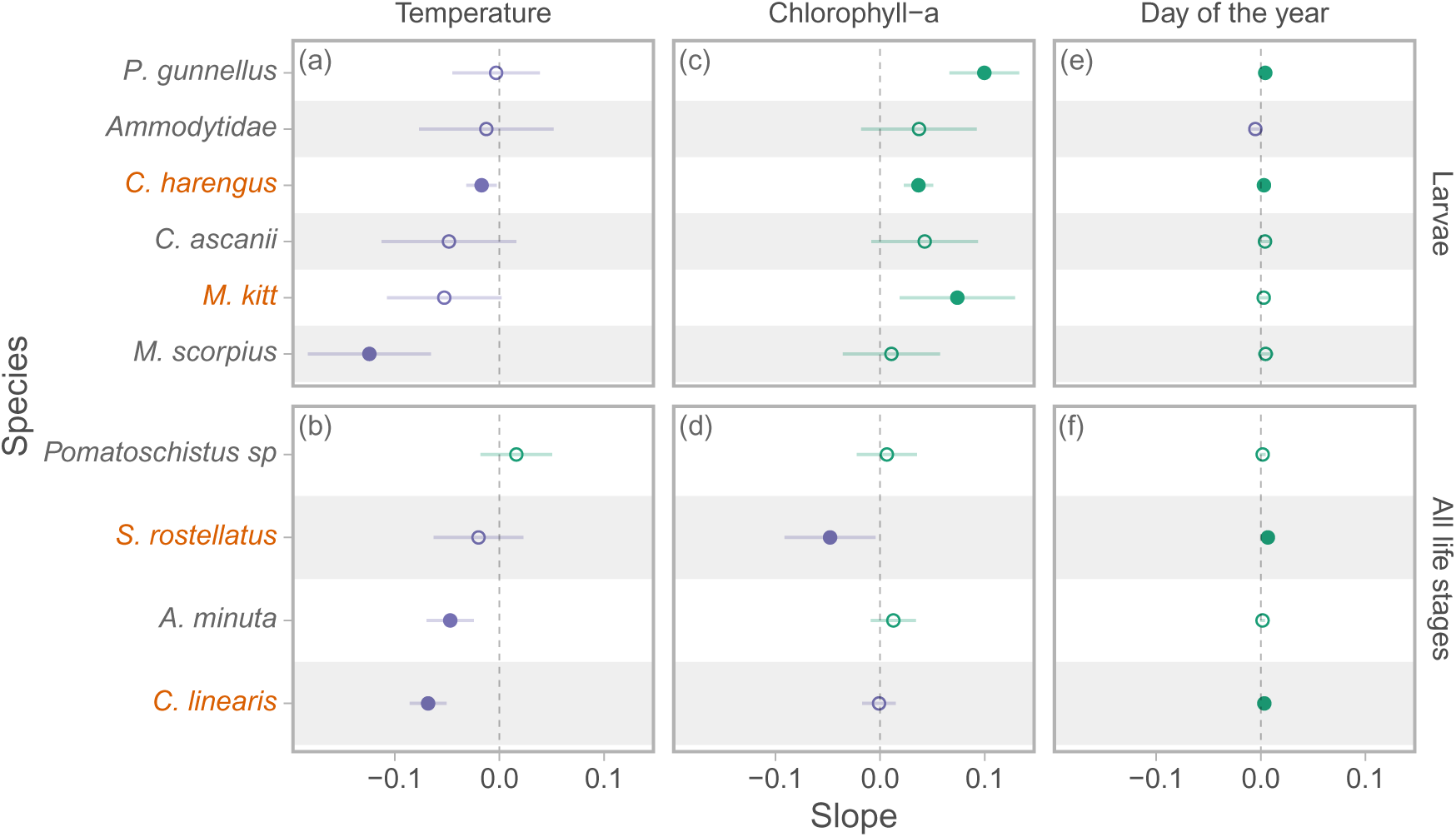
Estimated coefficients for temperature, chlorophyll-a, day of the year from the spatial generalized gamma model fitted to larvae length, by species. Purple points denote negative effects, green positive, and open points indicate that the 95% confidence interval of the estimate overlaps 0. Species predominantly found in Skagerrak are in orange text.

Spatial predictions of larvae lengths averaged over time reveal a clear tendency for a separation near the Skagerrak and Kattegat border (Figs. 2, 4), such that the sizes are generally larger in Kattegat than in Skagerrak. The only two exceptions to this are *Pomatoschistus* spp. and *S. rosellatus*, which to some degree show an opposite pattern (Fig. 4). The overall spatial pattern is fairly consistent over time, see e.g., Supporting Information Figs. S7, S8 for spatial predictions by year for two selected species (*Ammodytidae* and *Pomatoschistus* spp). The average spatial patterns in density from the spatiotemporal models did not show the same similarity across species as in the length models (Supporting Information Fig. S9). Also in the density models, the spatial patterns are relatively consistent over time, see e.g., Supporting Information Figs. S10, S11 for predictions by year for two selected species (*Clupea harengus* and *Pholis gunnellus*). While the abundance indices of *Clupea harengus* especially had declined drastically in the time period, species such as *Ammodytidae* have increased slightly. For some species (*Chirolophis ascanii*, *Myoxocephalus scorpius*, and *Agonus cataphractus*), the center of gravity showed very little variation over time and varied less than 12 km in any direction over 30 years (Fig. S2). *Aphia minuta* and *Pomatoschistus* spp. showed the strongest temporal trends in center of gravity, with average latitudes increasing by 24 and 14 km per decade, respectively. The remaining species showed intermediate annual variation (X and Y coordinates varying less than 10 and 50 km over time) without trends (Fig. S2).

**Figure 4:**
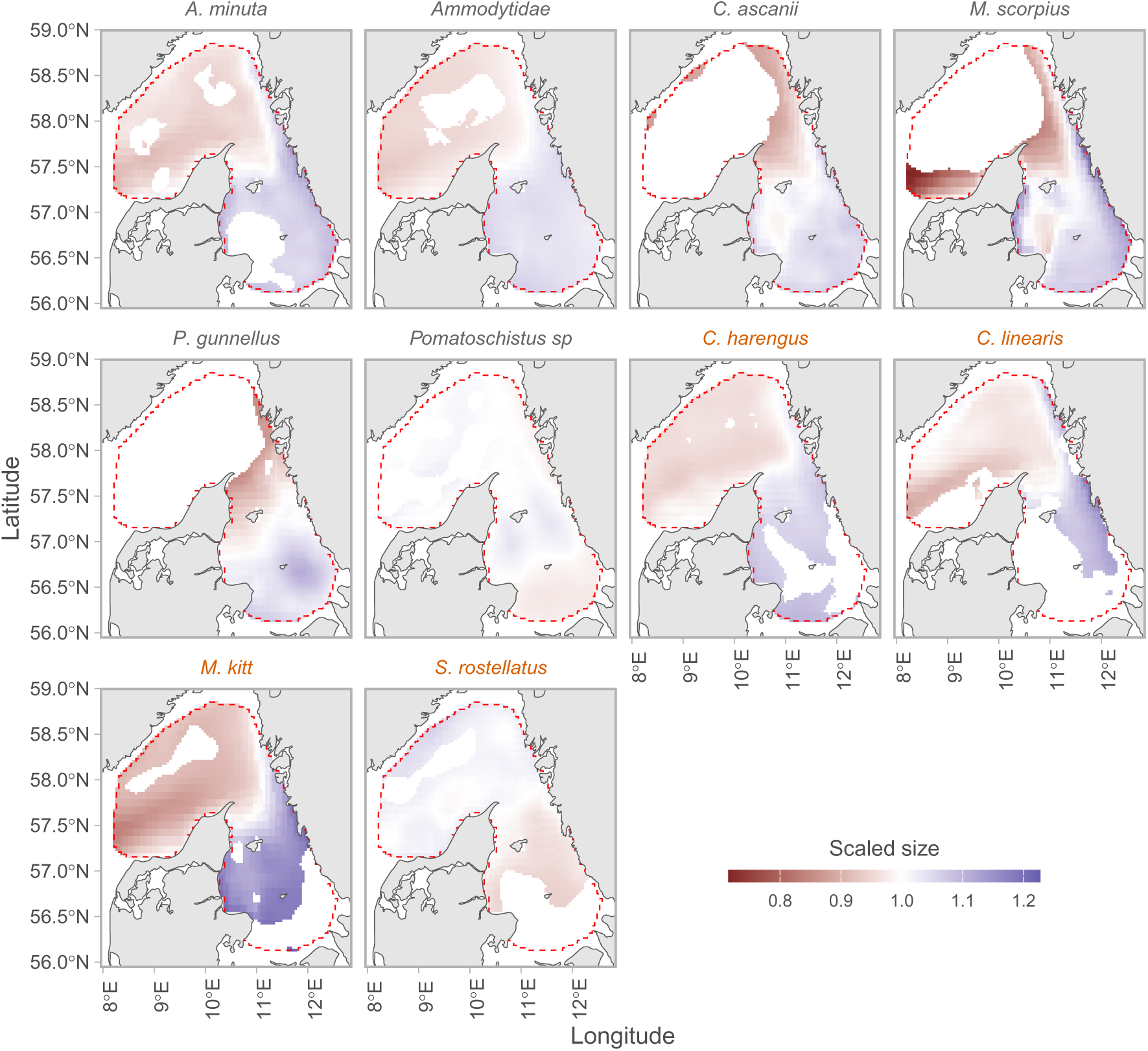
Spatial patterns of predicted average larvae lengths, normalized to the geometric mean for each species to allow for comparison, such that red areas correspond to lower than average sizes and blue correspond to large than average sizes. The red dashed line encompasses the spatial domain. Only grid cells that cumulatively make up 95% of the density for the species are included. Species predominantly found in Skagerrak are in orange text.

Since around 2010, the January density of *Calanus* spp. declined in Skagerrak and Kattegat. However, comparable low densities where also observed in Kattegatt prior to this, in the 1990’s and early 2000’s. In Kattegat, a strong seasonal trend with peaks in March and September was observed. In contrast, the seasonal dynamics in Skagerrak were less clearly bimodal (Fig. 5, 6a, Supporting Information Fig. S13). The January density of large copepods showed tendencies of declines over time in Skagerrak, but less so in Kattegat. Highest densities are found in the summer months (Fig. 5, 6b).

**Figure 5:**
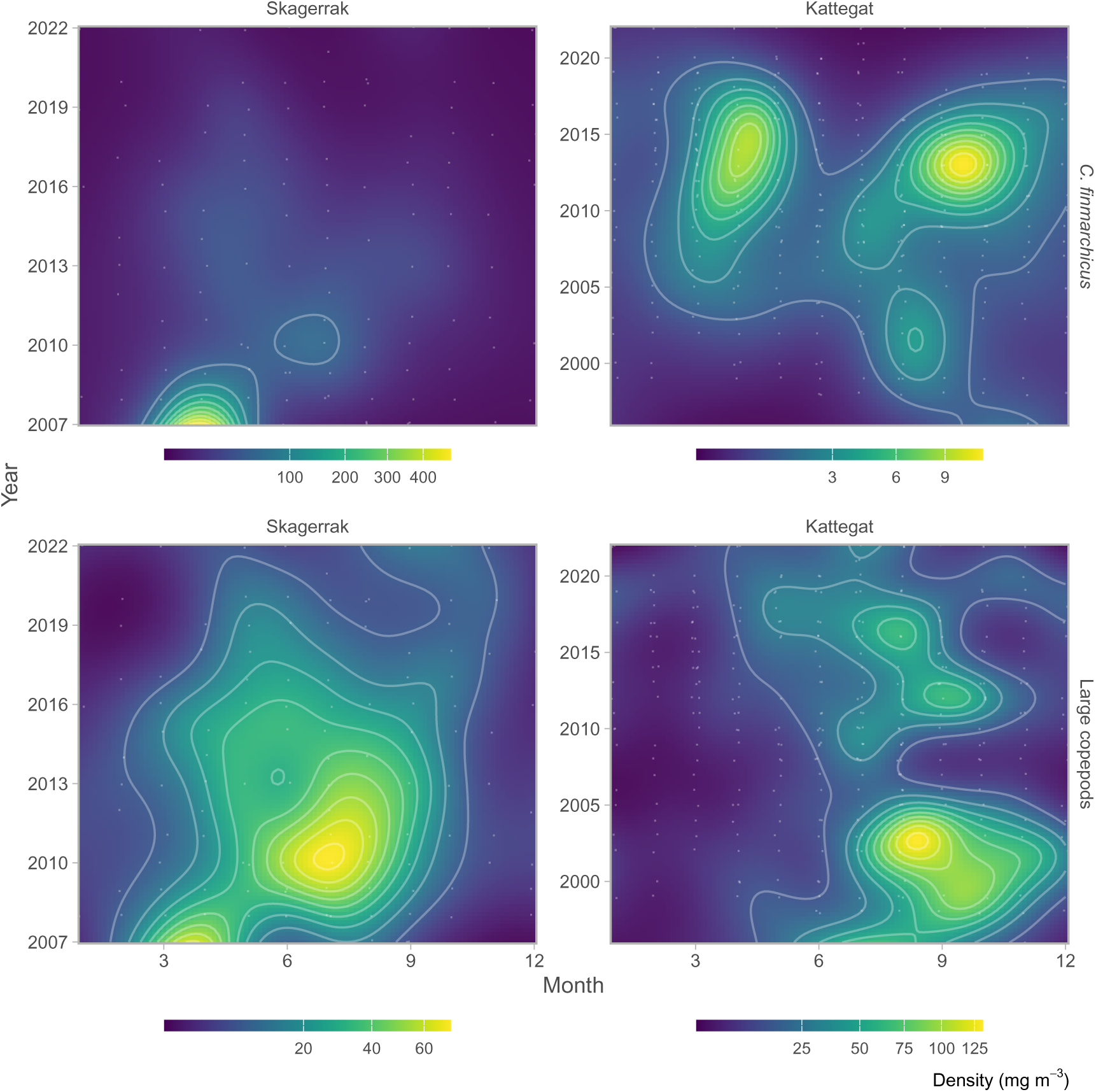
Seasonal and long term trends in the density of *Calanus* spp. and large copepods (*>*0.25 mm). The top row shows the predicted densities of *Calanus* spp. and the bottom row shows predicted large copepod density, for Skagerrak (left) and Kattegat (right), as a function of month (x-axis) and year (y-axis). Note the scales differ for each subplot.

The density-weighted standardized indices of mean larvae length based on predictions from the spatial model revealed declines in average sizes over time, especially since mid 2000’s. Since 2010, the percent change in size was on average −14% and −9% in Skagerrak and Kattegat, respectively, ranging between −30%– −1% across all species (Fig. 6d, 7). Most species also exhibited a substantial variation in length from year to year. Analysis of the global trend in fish size based on hierarchical GAMs fitted to *z*-scored time series revealed a clear decline since 2010 (Fig. 5, Supporting Information Fig. 6c). To some degree, the *Calanus* spp. time series co-vary with time series of length, especially in Skagerrak (Fig. 6a).

**Figure 6:**
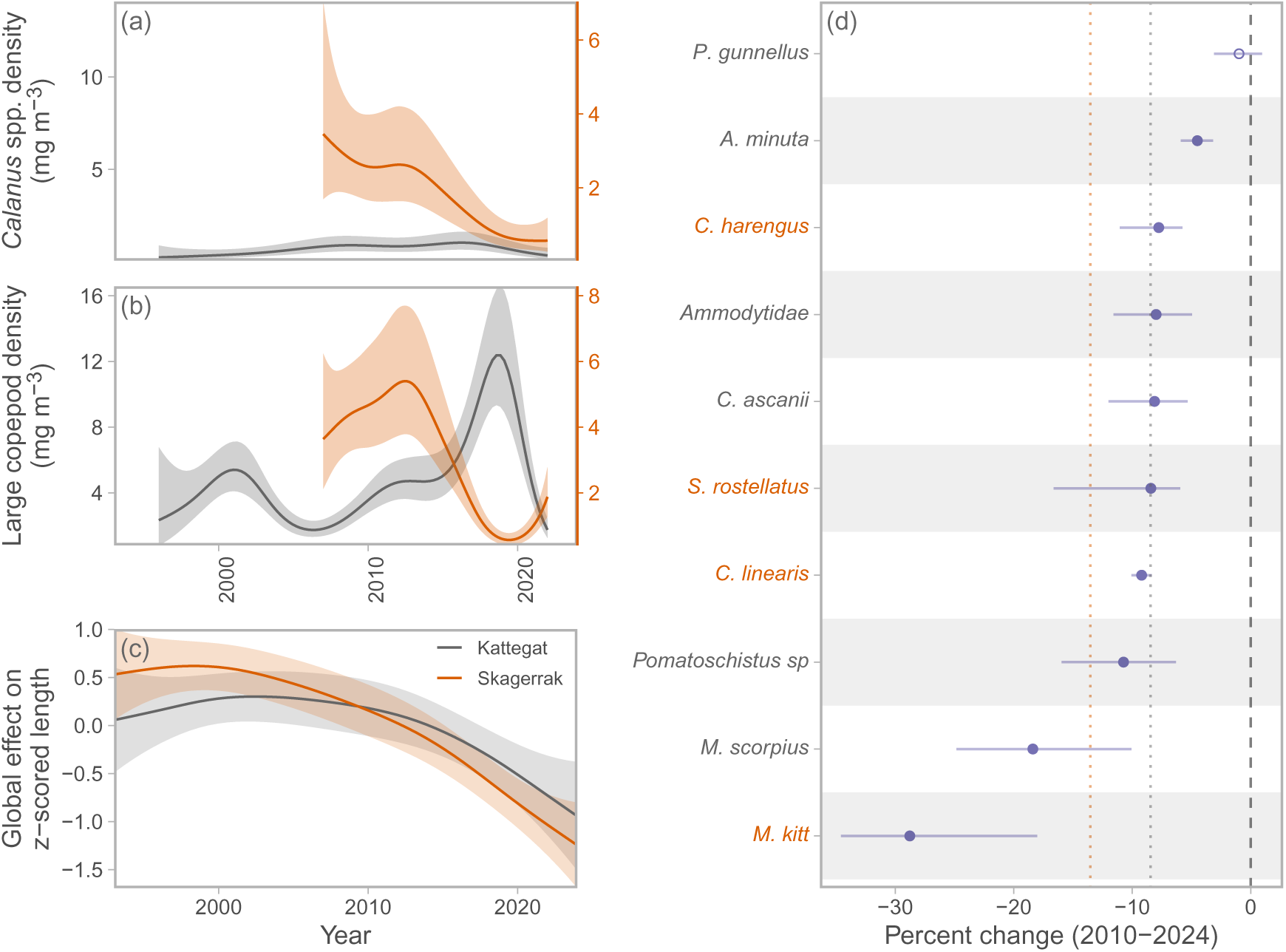
Trends in plankton densities and body lengths. Panel a) shows the predicted densities of *Calanus* spp. in January for Skagerrak (orange line and right y-axis) and Kattegat (grey line and left y-axis), panel b) shows the predicted density of large copepods, and c) shows the global prediction from the hierarchical GAM fitted to *z*-scored lengths. Panel d) shows the species-specific percent change in body size since 2000, which marks the peak in body length from the global prediction. In panel d), points indicate the median and the horizontal line covers the 10^th^–90^th^ percentile across 500 draws from the joint precision matrix. Filled circles indicate the the confidence interval does not overlap 0. Species predominantly found in Skagerrak are in orange text.

**Figure 7:**
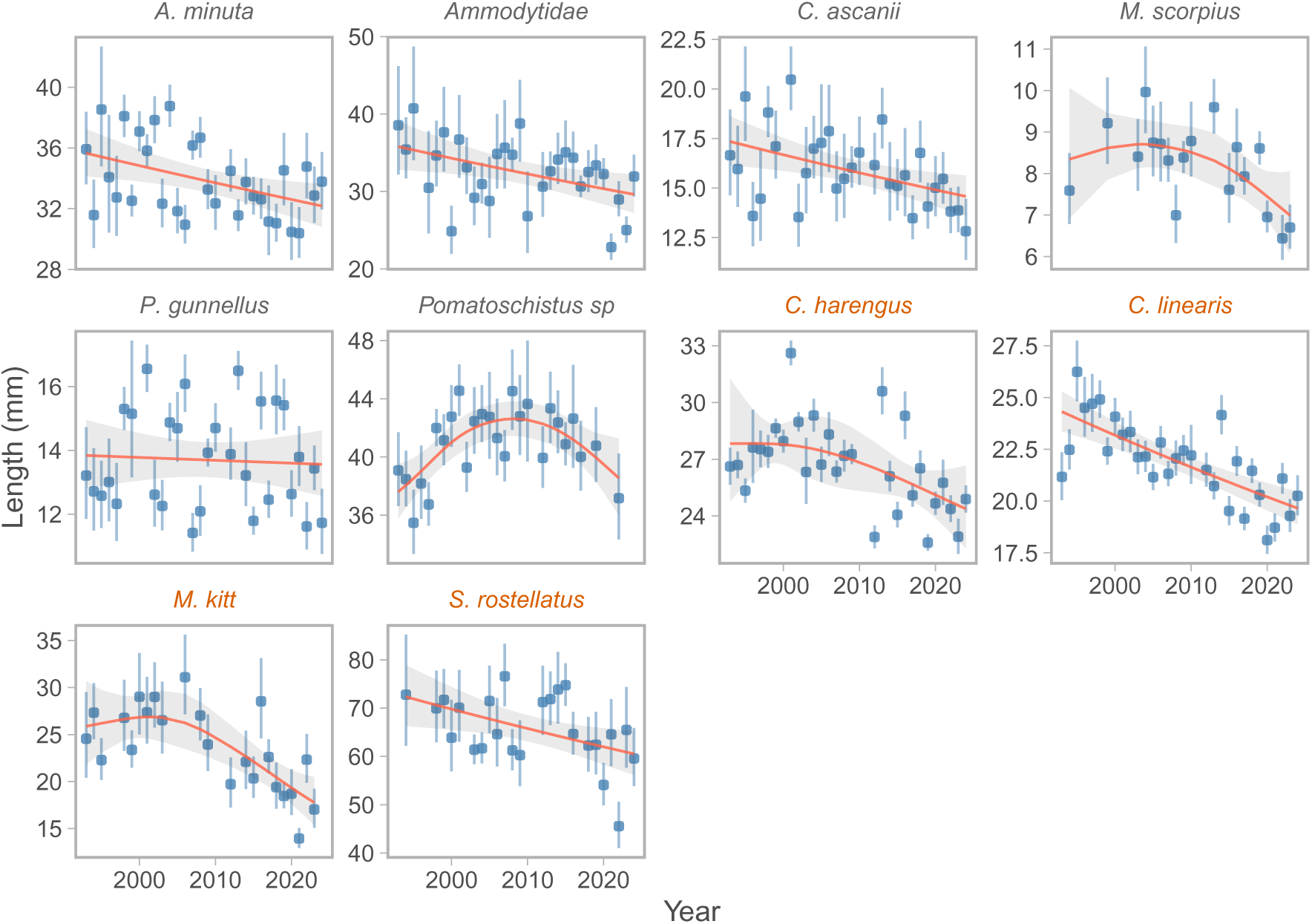
Trends in density-weighted indices of larvae length from the spatial generalized gamma model. Blue points correspond to annual indices and the vertical error bars cover the 95% confidence interval. The red line and gray ribbon show the fit and the 95% confidence interval of a log-link lognormal generalized additive model with year modelled as a smooth effect with basis dimensions (*k*) of eight to illustrate trends over time. Species predominantly found in Skagerrak are in orange text.

## Discussion

Our analysis of three decades of ichtyoplankton surveys in the Skagerrak and Kattegat revealed consistent declines in larval body size across multiple non-commercial and commercial fish species as well as the size of small bodied species, in particular in the last 15 years. The observed average decline in length of −14% and −9% across species in Skagerrak and Kattegat, with some species showing more substantial decreases (up to −31%), aligns with global trends of decreasing body size in marine organisms (Daufresne *et al*. 2009, Martins *et al*. 2023). The pattern of size reductions was particularly evident in larvae, where temperature had a negative effect on size in all species. This result aligns with the hypothesis that early life stages may indeed be more vulnerable to temperature changes due to their narrower thermal tolerance ranges (Rijnsdorp *et al*. 2009, Pörtner and Peck 2010).

While we can establish that 9/10 estimates of temperature effects are negative, and temperature increased in the spatial domain, it is difficult to ascertain the mechanism underlying the negative size trends over time, because temperature can act on body size through multiple interacting and non-exclusive mechanisms. Since larvae have limited ability to behaviorally thermoregulate due to their relatively restricted swimming capacity, and limited ability to store energy reserves (Pörtner and Peck 2010), ocean warming ought to lead to higher experienced temperatures. This leads to higher metabolic rates, and thereby feeding rates. Unless larvae are able to meet the higher metabolic demand with increased feeding rates, due to a limited food supply, growth rates will decline and mortality from starvation can increase. Some of our results suggest the drivers of negative trends in size may be related to food availability. Firstly, we find positive effects of chlorophyll-a, which we here use as a proxy for prey availability. Secondly, time-series of fish sizes tend to co-vary with year-to-year trends of January density of both *Calanus* spp. and large copepods. Similar conclusions where also drawn in a study on North Sea herring, where reductions in larvae growth rate (inferred from otolith increments), was linked to declines in recruitment (Payne *et al*. 2013). Hence, it seems plausible that food availability and direct- and indirect effects of temperature may have driven the changes in fish sizes.

Climate change may also have acted on fish sizes via the links to their prey. *C. finmarchicus*, one of the copepods included in our indices, is a cold water copepod (Rees 1957, Jaschnov 1970) that has declined in the Northeast Atlantic since the early 1960’s (Planque and Fromentin 1996), and has exhibited a poleward shift at a rate outpacing many terrestrial organisms as a consequence of ocean warming (Beaugrand *et al*. 2002, Parmesan and Yohe 2003, Chust *et al*. 2014). Future projections under high greenhouse gas emission scenarios suggest that *C. finmarchicus* densities could decrease by up to 50% by the end of the century in some regions of the North Atlantic (Grieve *et al*. 2017). As the *C. finmarchicus* included in our analysis are from the southern edge of their geographical distribution (Planque and Fromentin 1996), we expect climate change to continue to have large negative effects on their abundance. Being a critical link between microzooplankton, including ciliates, as well as phytoplankton such as diatoms and flagellates, and higher trophic levels, this decrease in abundance could have wide ecosystem impacts.

Because we do not know the age of the fishes, only their size, we cannot determine if temperature and food availability acts on growth or size-dependent survival. In terms of survival, small larvae may be more susceptible to starvation by having higher energy requirements per unit body mass. Mortality from predation can also shape the size distribution of larvae. A commonly held view is that faster growth leads to higher survival because they more quickly grow out of the predation window. While it has been supported in many laboratory studies (see review in Leggett and Deblois 1994), tests in natural systems are more rare (but see Pepin and Myers 1991). However, this expectation stems from a larvae-centric view and their ability to avoid predators (Leggett and Deblois 1994), while predator-prey interactions depend on both predator and prey sizes (Olson 1996, Andersen 2019). For some predators it may be more beneficial to target smaller prey while for others it may be larger prey, which ever maximize the cost to benefit ration (according to the optimal foraging theory) (Stephens and Krebs 1986). Therefore, without detailed knowledge about the predation field our fishes are exposed to, which depends on the size structure of the predator community over time, we likely cannot state that mortality rates always is higher for smaller sized fish. The observation that temperature acts in similar ways for multiple species of different sizes also suggests that the main effect of temperature is a direct one, and not on size-selective mortality since these fishes experience predation from different predators.

There are several limitations of our study that are difficult to isolate but can have impacts on our inference. The larvae data we work with may for a given species come from multiple distinct populations or spatial spawning units. If these populations vary in size in a given day, for example due to differences in spawning date, this might have influenced our temporal analysis if the relative abundance of said spawning units have changed over time. In the example of herring, Kattegat and Skagerrak is populated from a mix of autumn and winter spawning herring drifted from North Sea larvae (Corten 1986), and likely also local populations. Observational data on early larvae from the main spawning grounds in the North Sea suggest an increase in the relative contribution of Winter spawning components of the North Sea herring stock (ICES 2024). Hence, the decreasing larval size in our samples from the Kattegat-Skagerrak is also compatible with an increasing proportion of Winter components which are significantly smaller at the time of the survey, and may represent a confounding effect.

Our study addresses a significant knowledge gap by examining long-term changes in both commercial and non-commercial species, focusing on small-bodied fish and larvae, because they are relatively understudied in relation to commercially important fishes. The demonstrated sensitivity of early life stages highlights that more research should focus on integrating these findings into population and food web models to better predict stock responses to various climate change scenarios. Enhanced predictions of early life stages growth in response to temperature and trends in prey availability could directly improve recruitment forecasts used in stock assessment and fishery advice which currently rely on simple statistical averages. Additionally, our results emphasize the need for management approaches that consider not only adult populations but also the environmental conditions influencing early life stages, which may become increasingly important determinants of population trajectories as climate change progresses. The consistent size reductions observed across different species suggests a generalized response to warming that could fundamentally alter marine community structure and function in the coming decades.

## Acknowledgements

We thank the personnel at the former Swedish Board of Fisheries, and at the current Institute of Marine Research, Department of Aquatic Resources, Swedish University of Agricultural Sciences, for long-time high-quality data collection during the Swedish part of IBTS.

## Author contributions

**ML:** Conceptualization (lead); Data Curation (equal); Methodology (lead); Formal analysis (lead); Writing – original draft (lead); Writing – review & editing: (lead) **MW:** Conceptualization (supporting); Data Curation (equal); Writing – original draft (supporting); Writing - review & editing (supporting) **PT:** Conceptualization (supporting); Data Curation (equal); Writing – original draft (supporting); Writing - review & editing (supporting) **FM:** Conceptualization (supporting); Methodology (supporting); Formal analysis (supporting); Writing – original draft (supporting); Writing - review & editing (supporting) **EQ:** Conceptualization (supporting); Data Curation (supporting); Formal analysis (supporting); Writing – original draft (supporting); Writing - review & editing (supporting) **VB:** Conceptualization (supporting); Writing – original draft (supporting); Writing - review & editing (supporting) **PJ:** Conceptualization (supporting); Writing – original draft (supporting); Writing - review & editing (supporting)

## Data Availability Statement

Code and data to reproduce the results are available on GitHub (https://github.com/maxlindmark/larvae-sizes/) and will be deposited on Zenodo upon publication.

## Funding

M.L. was supported by a research grant from the Swedish Research Council Formas (grant no. 2022-01511 to Max Lindmark).

## Notes

https://github.com/maxlindmark/larvae-sizes

### Competing Interest Statement

The authors have declared no competing interest.

