## Supporting Information for "Reduced size of larvae and small fish linked to warming and reduced prey density"

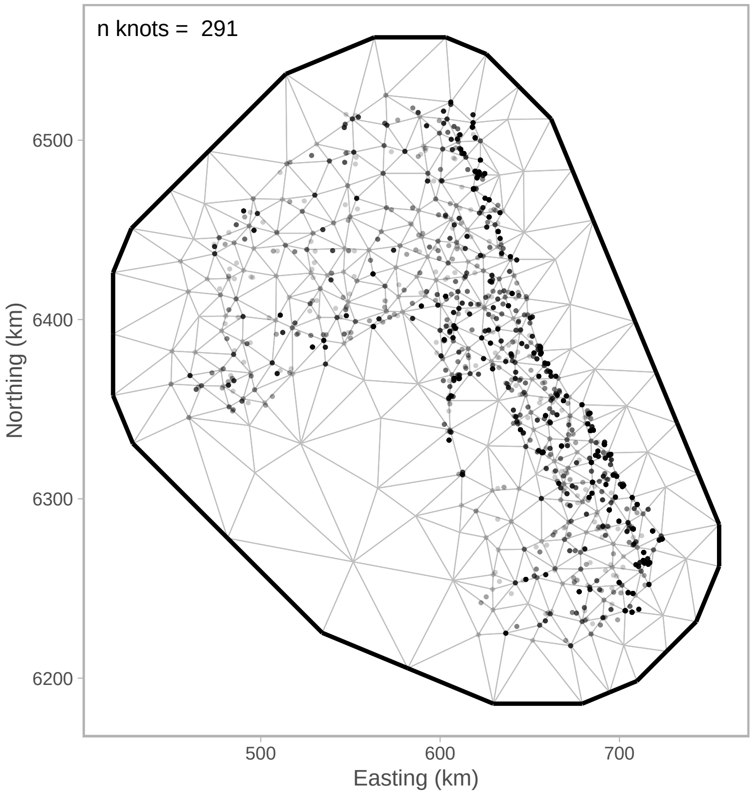


Figure S1: Example SPDE mesh (*Aphia minuta*).

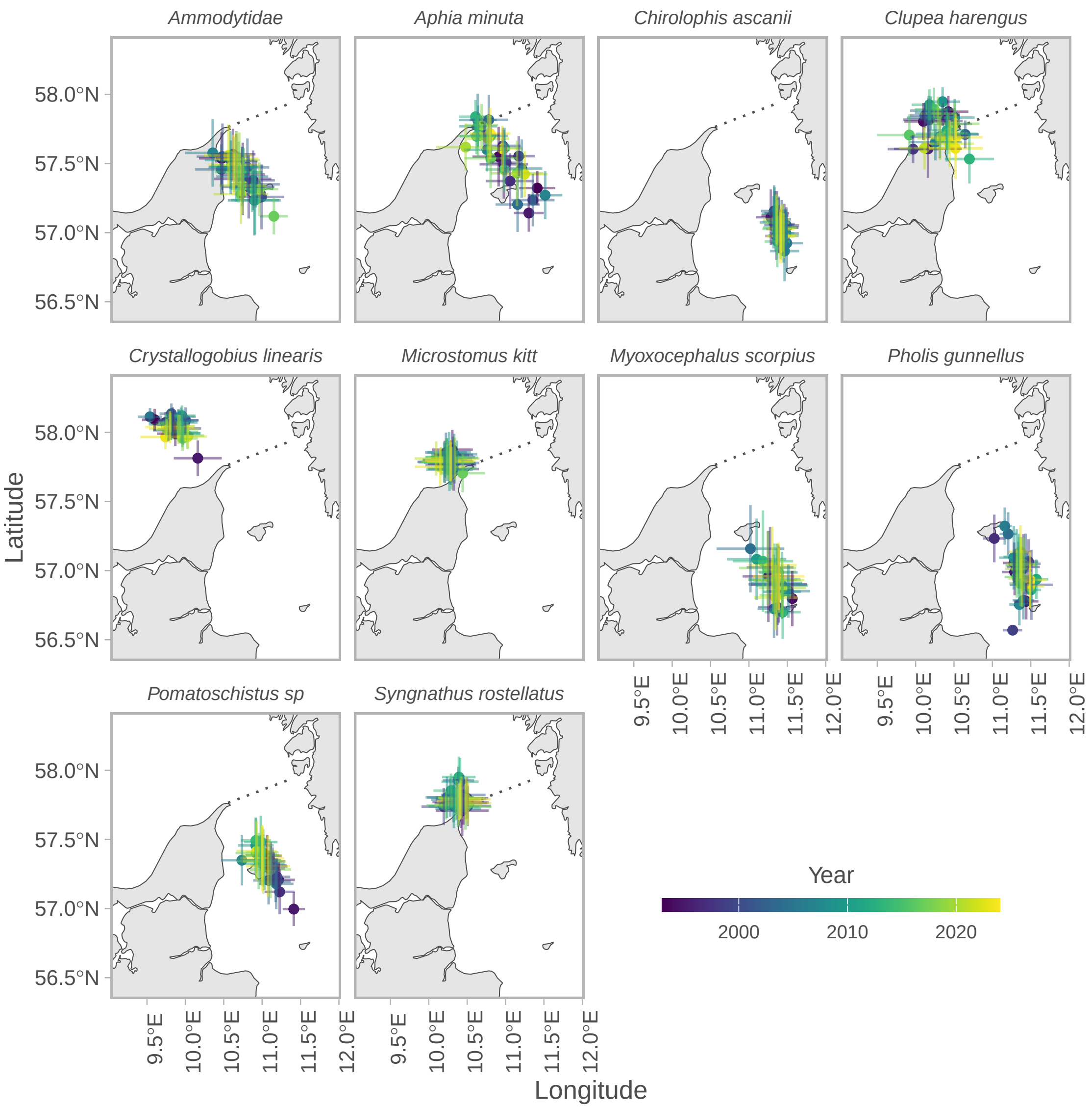


Figure S2: Center of gravity in X and Y coordinates (points) and their associated 95% confidence intervals, derived from the spatiotemporal density models.

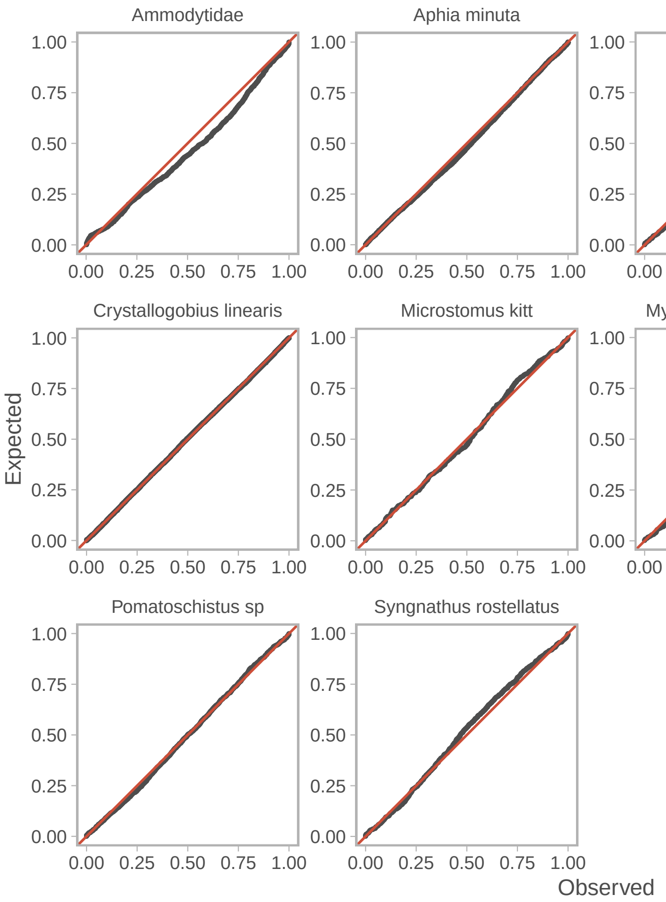


Figure S3: QQ-plots of the larvae length models based on simulated randomized quantile residuals (Dunn *et al.* 1996, Hartig 2022).

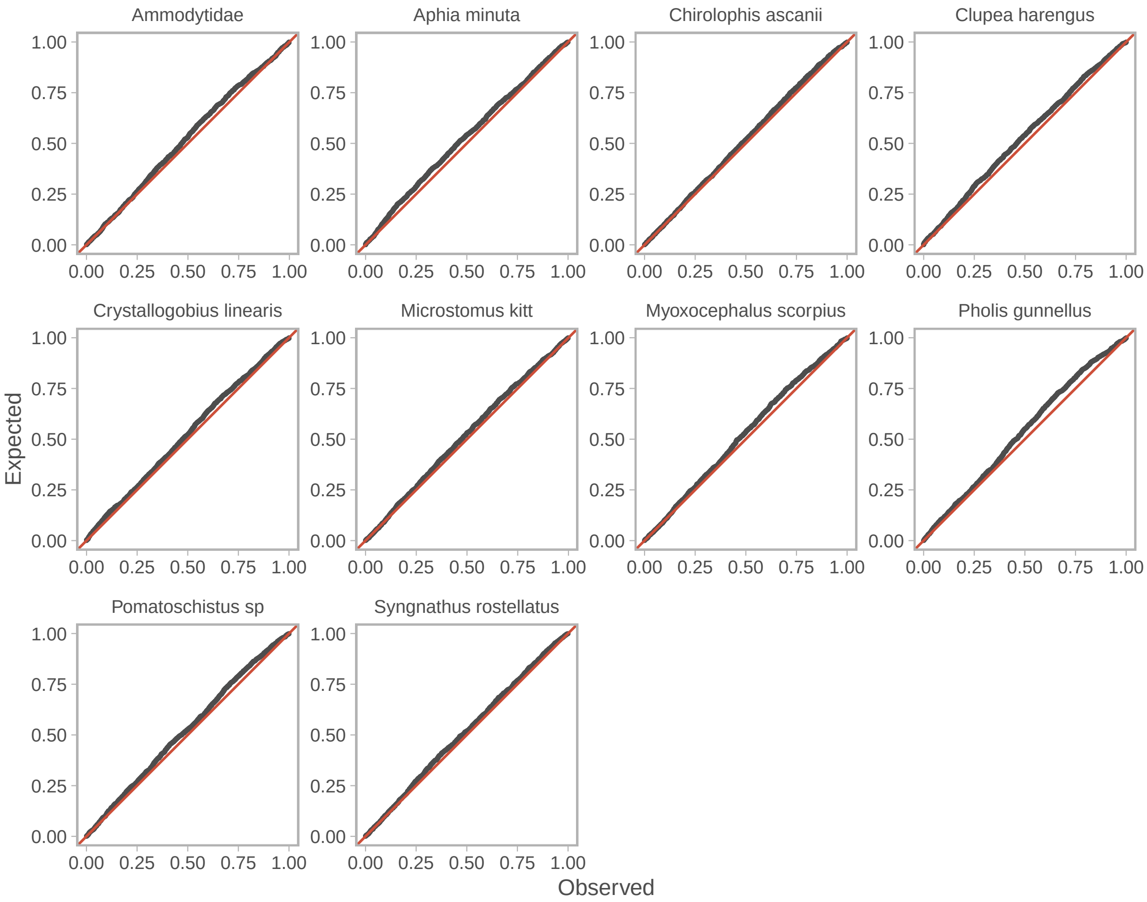


Figure S4: QQ-plots of the larvae density models based on simulated randomized quantile residuals (Dunn *et al.* 1996, Hartig 2022).

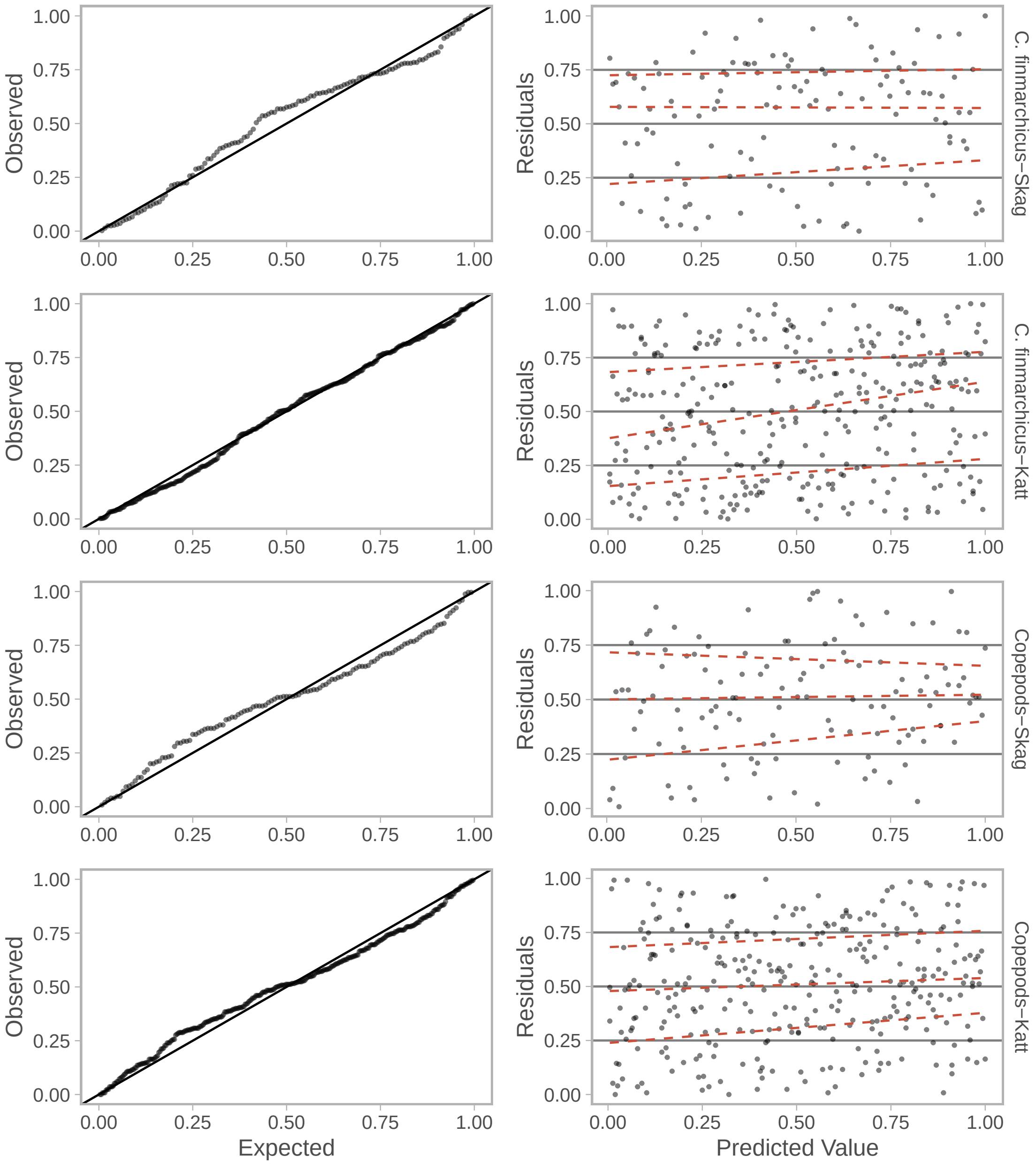


Figure S5: QQ-plot and residuals against the predicted value for the GAMs fitted to *Calanus* spp. and large copepod densities in Skagerrak and Kattegatt.

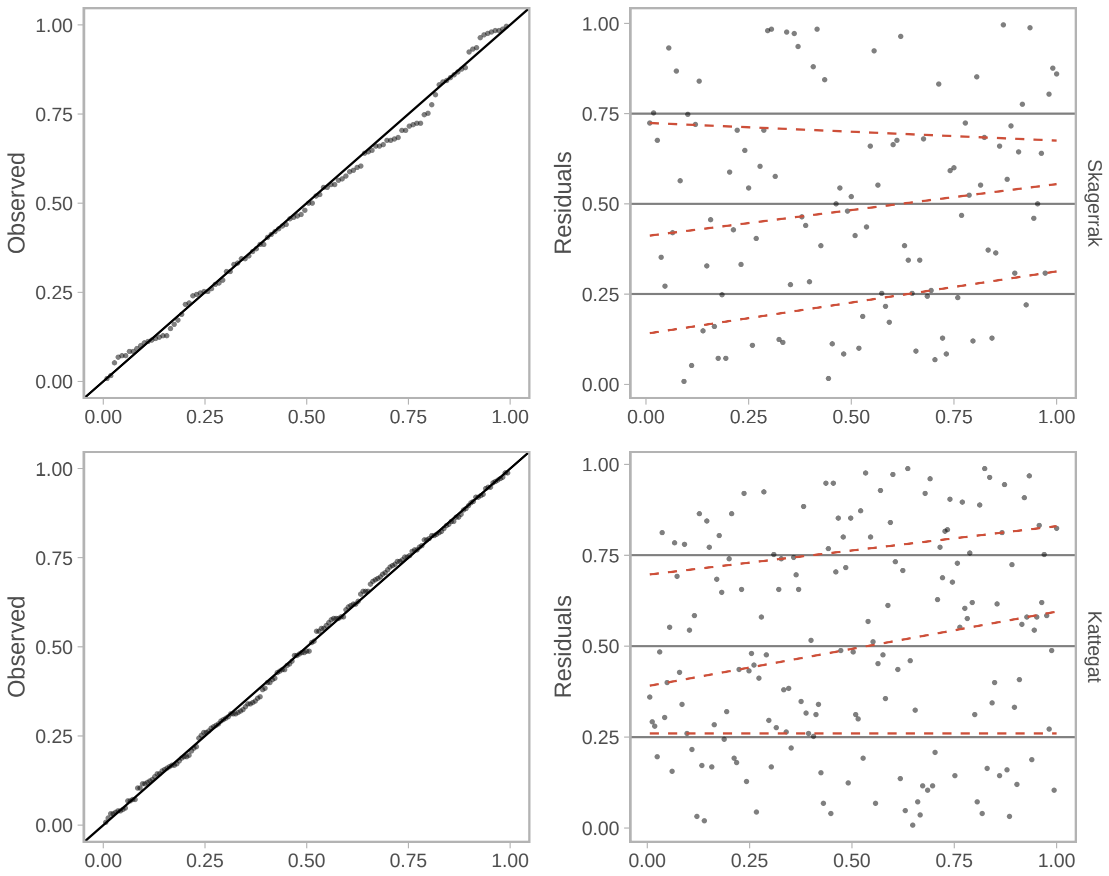


Figure S6: QQ-plot and residuals against the predicted value for the hierarchical GAM fitted to *z*-scored body lengths for Skagerrak (top) and Kattegat (bottom).

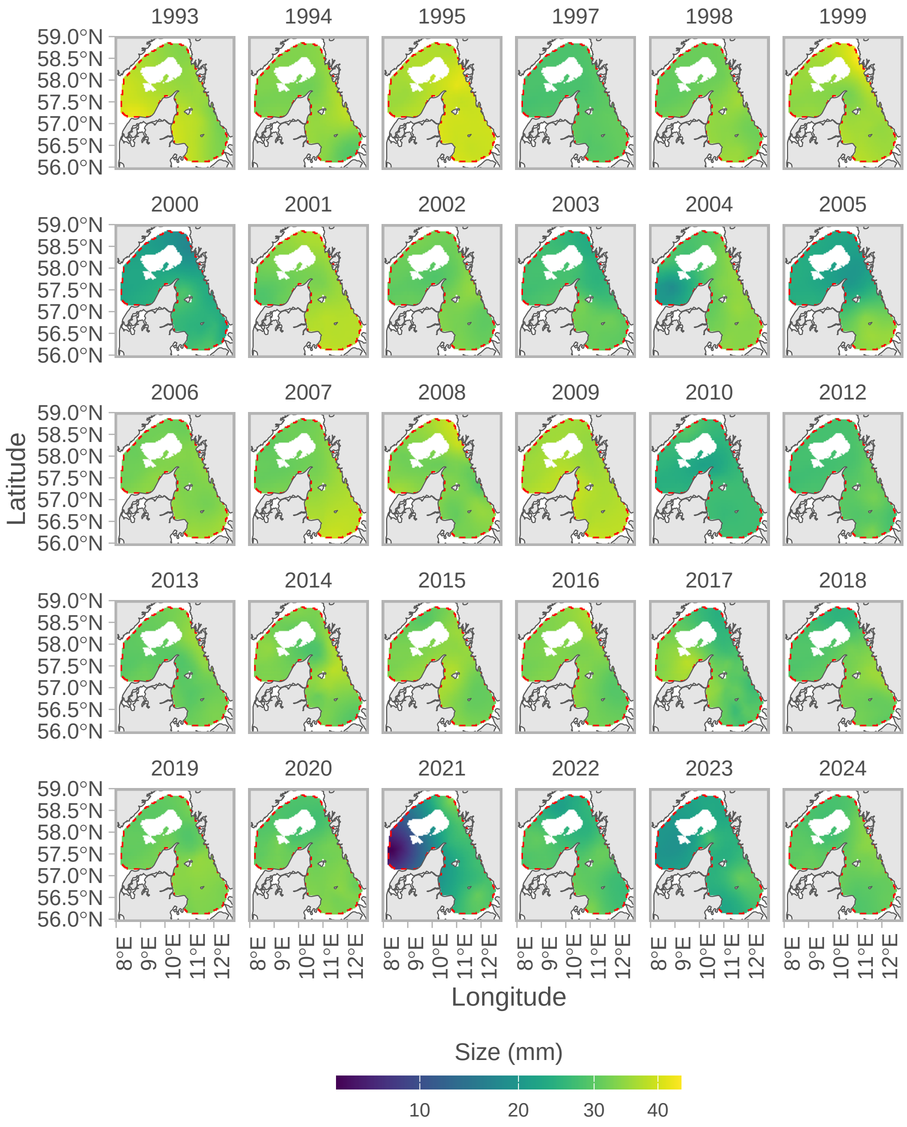


Figure S7: Predicted length (mm) over time in space for *Ammodytidae*. The color scale is square-root transformed to better visualize spatial patterns.

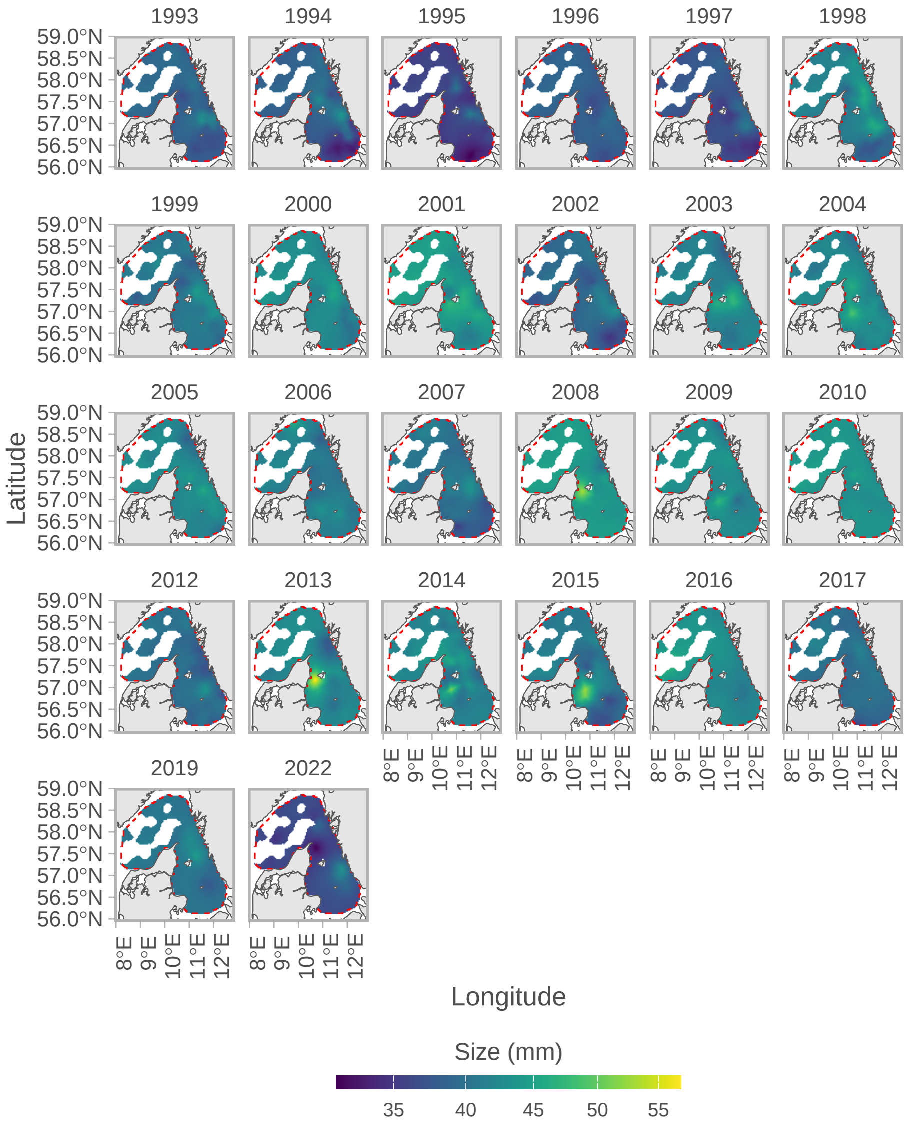


Figure S8: Predicted length (mm) over time in space for *Pomatoschistus* spp. The color scale is square-root transformed to better visualize spatial patterns.

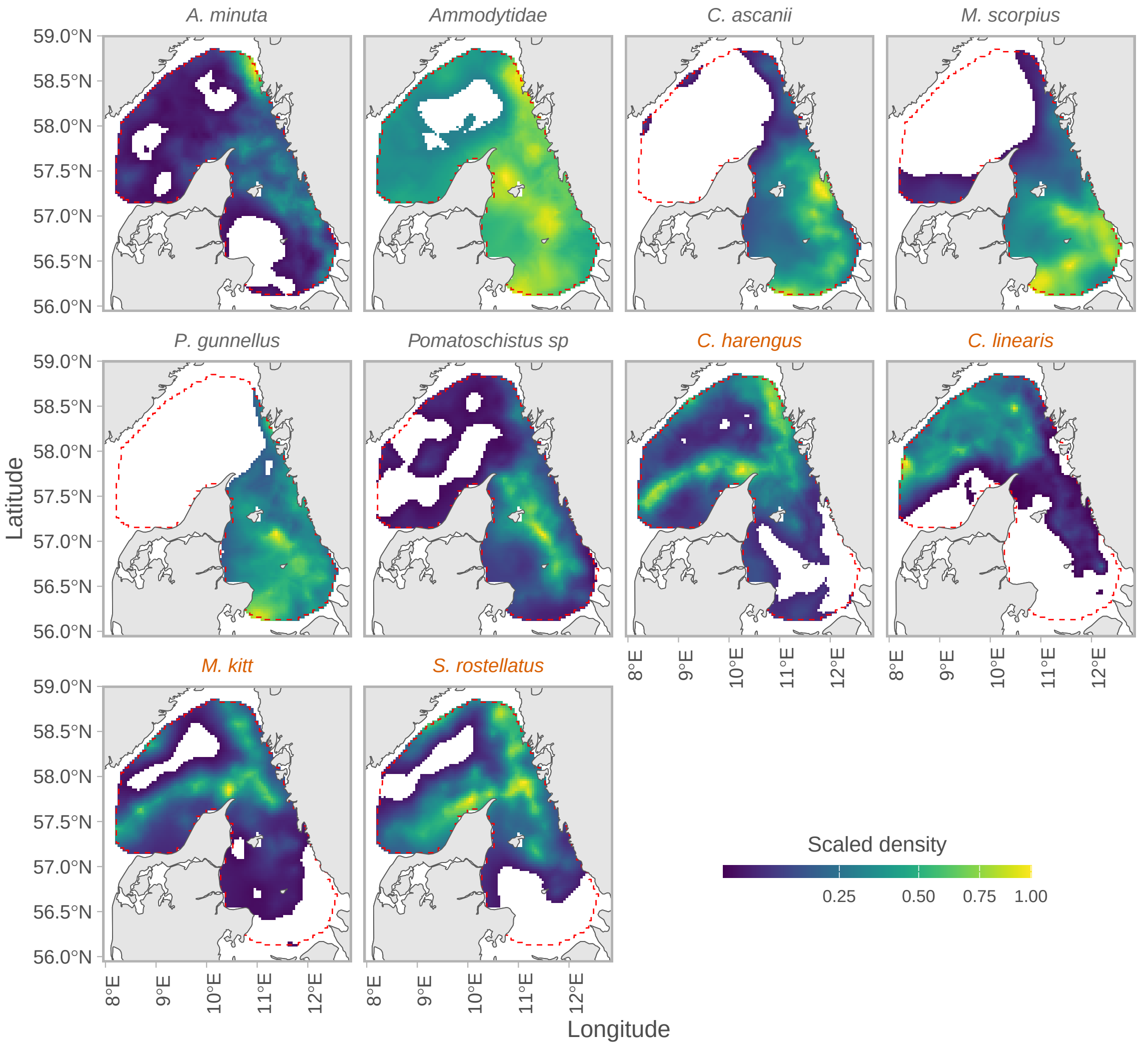


Figure S9: Spatial patterns of predicted average larvae density, normalized to the mean for each species to allow for comparison, such that blue areas correspond to lower than average densities and yellow correspond to large than average densities. Note the color scale is square root transformed to better visualize patterns. Species predominantly found in Skagerrak are in orange text.

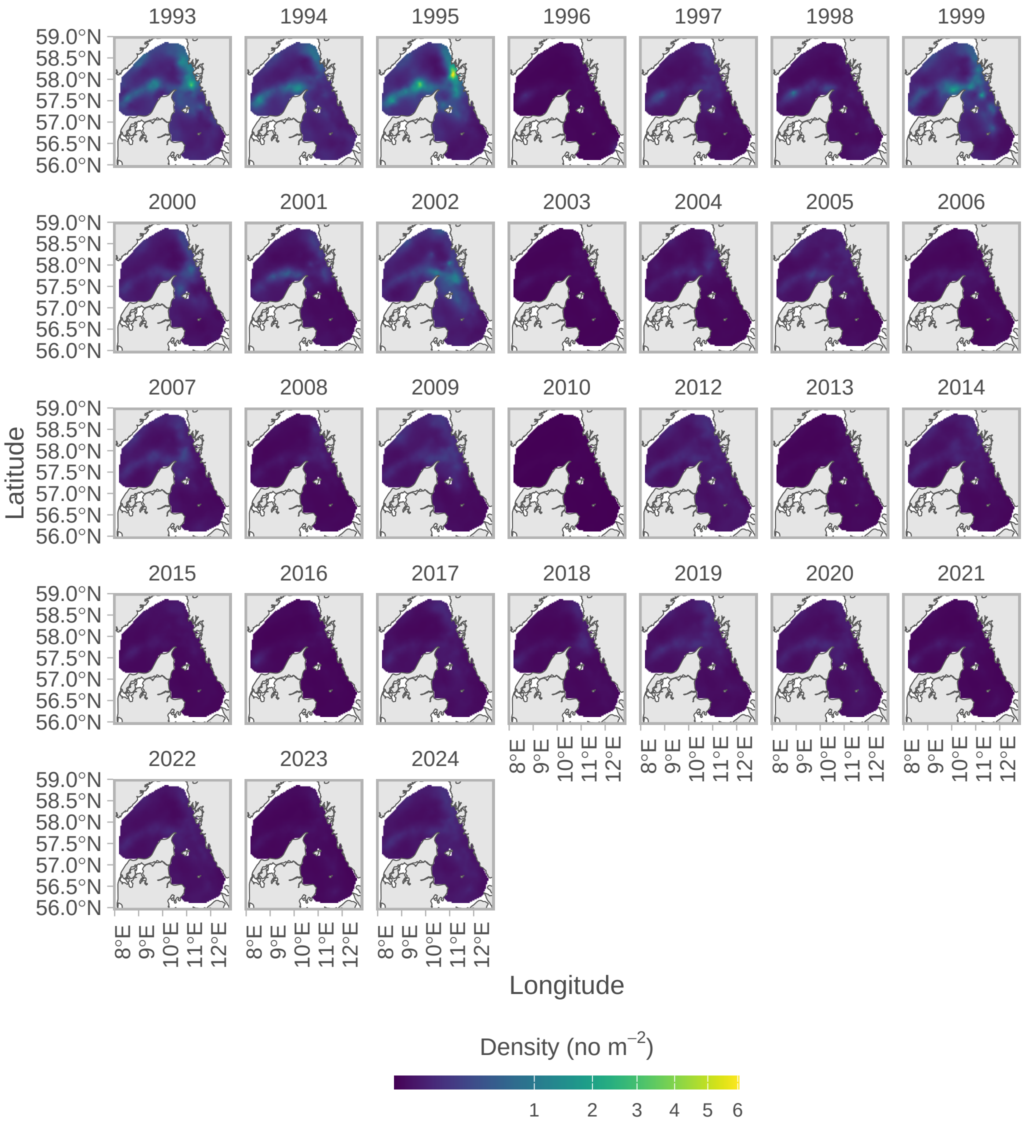


Figure S10: Predicted abundance density (no m$^{-}$2) over time in space for *Clupea harengus*. The color scale is square-root transformed to better visualize spatial patterns.

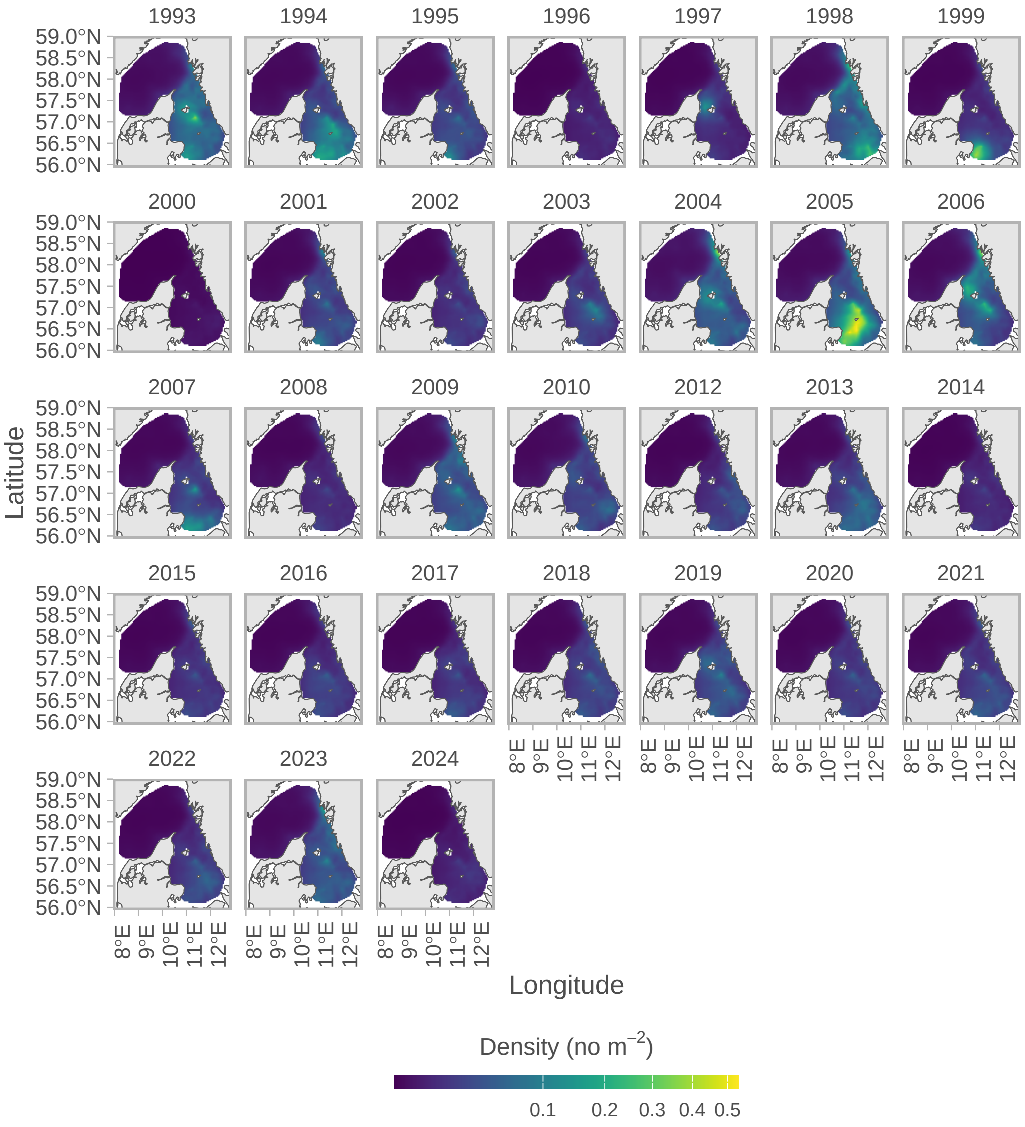


Figure S11: Predicted abundance density (no m$^{-}$2) over time in space for *Pholis gunnellus*. The color scale is square-root transformed to better visualize spatial patterns.

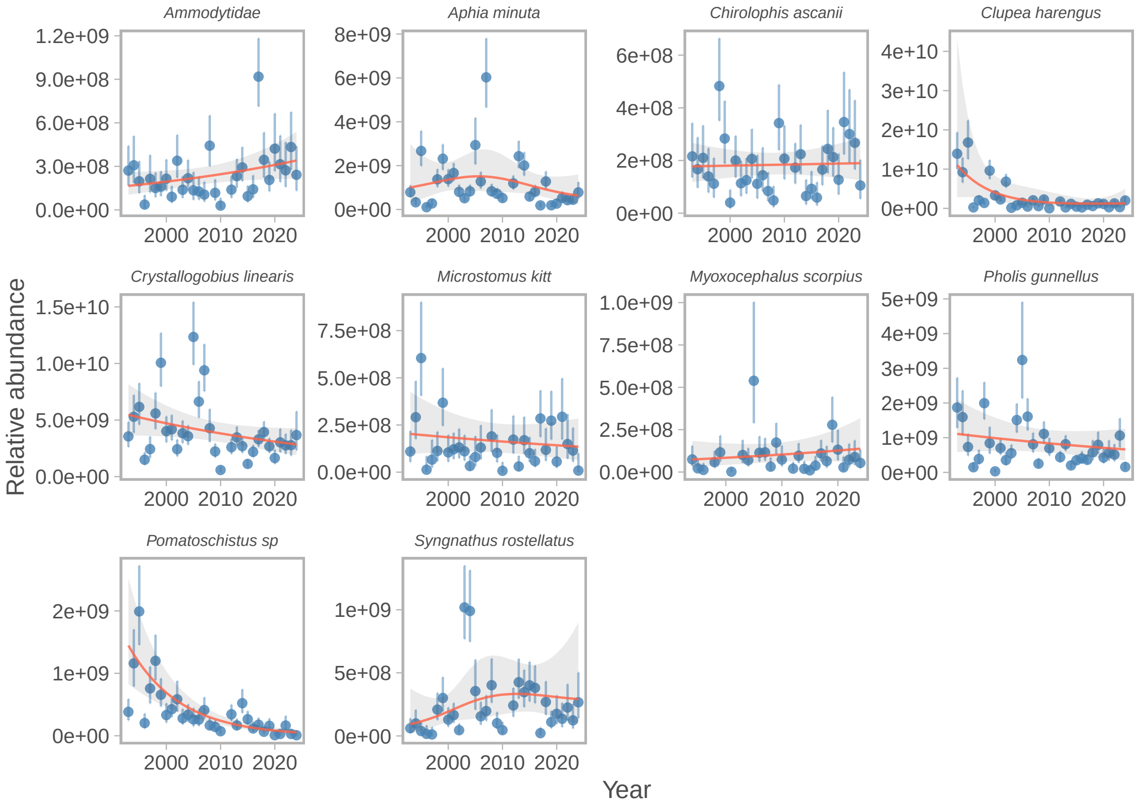


Figure S12: Trends in relative larvae abundance from the spatiotemporal, poisson-link delta-gamma model. Blue points correspond to annual index and the vertical error bars cover the 95% confidence interval. The red line and grey ribbon show the fit and the 95% confidence interval of a log-link lognormal generalized additive model with year modelled as a smooth effect with basis dimensions ($k$) of eight to illustrate trends over time.

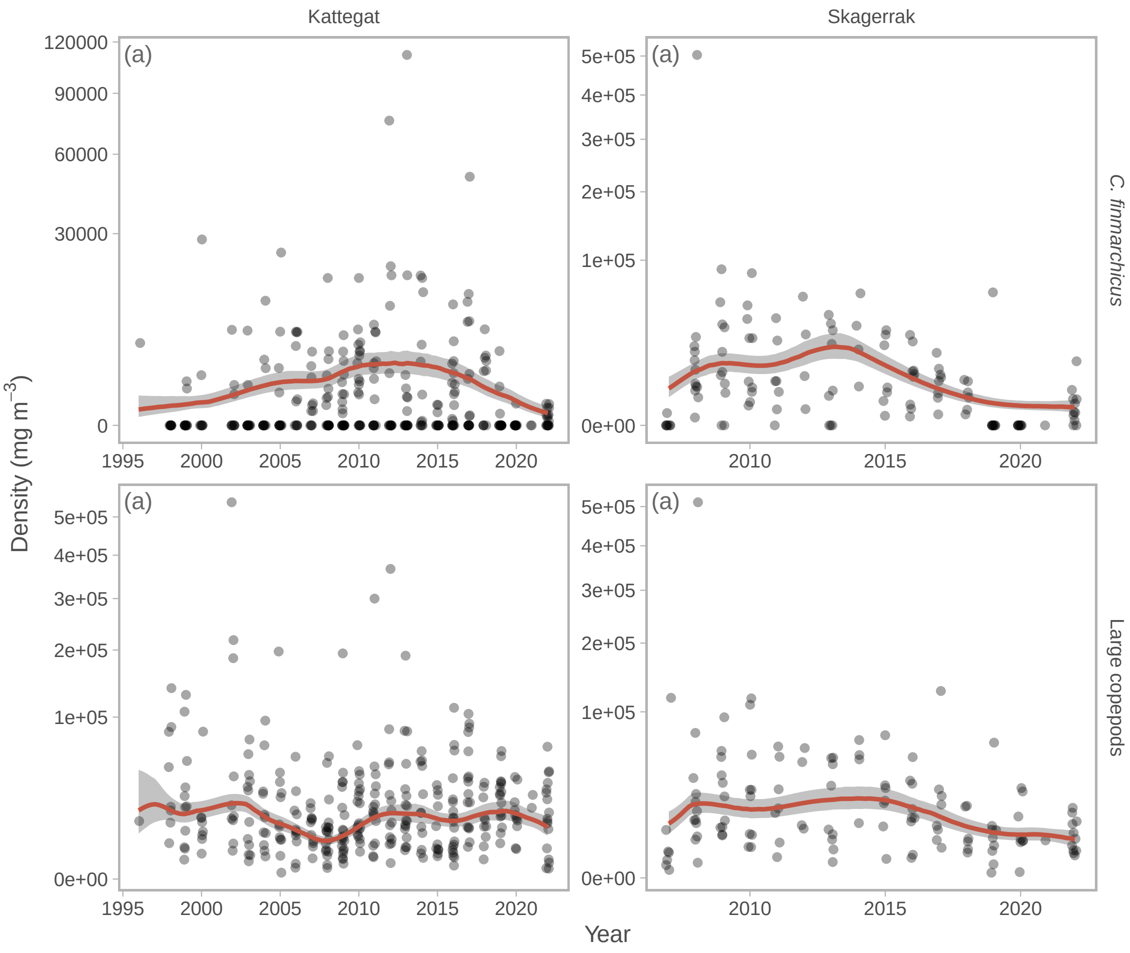


Figure S13: Predicted (lines) vs observed (points) *Calanus* spp. (top) and large copepod densities (bottom), for Kattegat (left) and Skagerrak (right). Predicted lines are averaged by year, while each data point is a single sample.

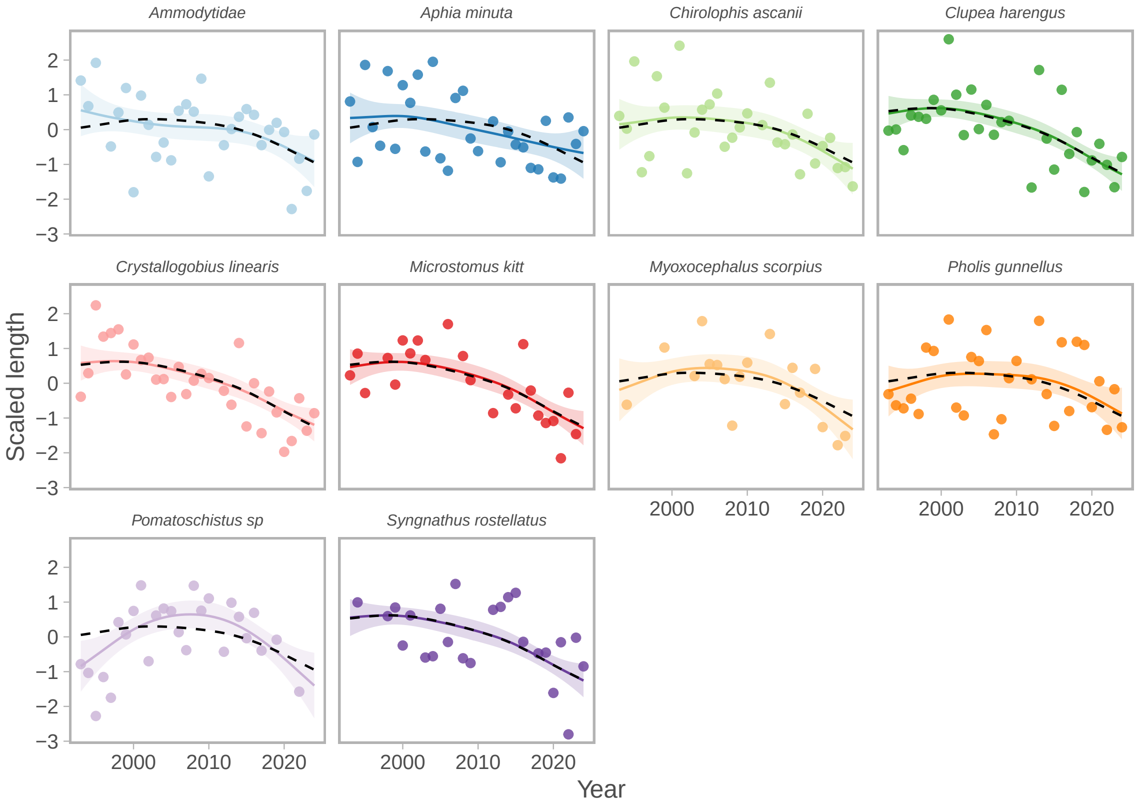


Figure S14: Trends $z$-scored length for each species and predictions from the hierarchical GAM. Coloured lines show the species-specific prediction, and the black dashed line the global prediction.
